# Microbiome-mediated Biotransformation of Human Lactoferrin (effera^®^) Enhances Epithelial Barrier Function in an Ex Vivo Adult Model

**DOI:** 10.64898/2026.08.03.742593

**Authors:** Nicole Kaplan, Rajitha Gadde, Ross Peterson, Pieter Van den Abbeele, Anthony Clark

## Abstract

Lactoferrin is a multifunctional iron-binding glycoprotein that supports intestinal barrier function, immune regulation, and a favorable gut microbial environment. However, the contribution of gut microbial biotransformation to its gastrointestinal activity remains poorly understood. We investigated whether effera^®^, a precision fermentation-derived recombinant human lactoferrin, supports intestinal barrier function through microbiome-mediated mechanisms. effera^®^ underwent simulated upper gastrointestinal digestion followed by ex vivo colonic fermentation using the validated SIFR® technology pipeline, which employs bioreactors that are inoculated with fecal microbiota from six healthy adult donors. Microbial activity was evaluated by measuring short-chain fatty acid (SCFA) production, bacterial cell density, and microbiome composition. effera^®^ produced dose-dependent increases in the production of SCFAs and bacterial cell density demonstrating enhanced microbial metabolic activity. These metabolic changes were accompanied by shifts in key microbial groups within Bacillota_A and Bacteroidota. Intact effera^®^ and cell-free post-colonic fermentation-derived products were evaluated in a Caco-2/THP-1 epithelial-immune co-culture model under basal and lipopolysaccharide-challenged conditions. Whereas intact protein did not significantly improve epithelial barrier integrity, effera^®^’s post-colonic fermentation-derived products significantly enhanced transepithelial electrical resistance (TEER) under basal conditions and produced an even stronger barrier-protective response following LPS challenge. Across matched doses, effera^®^ consistently generated greater TEER responses than bovine lactoferrin. Improved barrier function was accompanied by increased expression of tight-junction-associated targets ZO-1 and occludin and reduced secretion of CXCL-10 and IL-8. Together, these findings demonstrate that microbial biotransformation enhances the biological activity of effera^®^, linking increased microbial metabolism with improved epithelial barrier integrity and modulation of inflammatory signaling. This integrated study provides a strong mechanistic foundation for the use of human lactoferrin in adult gut-health applications and offers valuable guidance for future adult clinical studies and infant-relevant investigations.

## Introduction

The gastrointestinal tract serves as a critical biological interface linking nutrition, microbial metabolism, and immune signaling. It is the primary site for nutrient digestion and absorption,^1^ and transport into the blood or lymphatic system. In addition, the gut functions as a selective barrier that prevents the translocation of harmful substances and pathogens into the circulation, thereby reducing the risk of inflammation.^2,3^ The intestinal epithelium participates in immune regulation through the secretion of cytokines, chemokines, and antimicrobial peptides.^4–7^ As a result, maintaining intestinal health and barrier integrity is essential for overall physiological well-being.

A key factor regulating intestinal health is the vast community of microorganisms residing in the gut, collectively referred to as the gut microbiome or microflora.^8^ The human gastrointestinal tract harbors approximately 100 trillion (10^14^) commensal microorganisms, forming a highly complex and metabolically active ecosystem.^9,10^ Growing evidence suggests that gut microbial composition and metabolite production influence physiological processes extending far beyond digestion, including inflammatory regulation, energy metabolism, mitochondrial function, and overall host health.^11,12^ One of the primary metabolic outputs of gut bacteria is the production of short-chain fatty acids (SCFAs),^13^ such as acetate, propionate, butyrate, and valerate.^14^ SCFAs exert many of their beneficial effects by preserving intestinal barrier integrity,^15^ suppressing inflammation, modulating the immune system,^16^ and supporting cellular energy metabolism.^17^ These metabolites enhance the expression of tight junction (TJ) proteins^18^ and stimulate mucus production, thereby strengthening the epithelial barrier.^14^ Butyrate has been shown to upregulate TJ proteins such as claudin, zonula occludens-1 (ZO-1), and occludin, which regulate paracellular permeability.^19^ Furthermore, butyrate mitigates lipopolysaccharide (LPS)-induced barrier dysfunction.^20^ Propionate promotes intestinal goblet cell differentiation and increases mucus production.^21^ Beyond their effects on barrier function and maintaining barrier integrity, SCFAs influence leukocyte chemotaxis, appetite regulation, lipid metabolism, blood lipid levels, cytochrome P450 enzyme maturation, insulin sensitivity, and glucose homeostasis, highlighting their broad role in maintaining health.^22–25^

In addition to SCFA production, gut microbial bacteria contribute to protein and amino acid metabolism, generating compounds such as histamine,^26^ γ-aminobutyric acid (GABA),^27^ and other signaling molecules.^28^ They aid in synthesis of the essential nutrient vitamin K,^29^ metabolize bile acids,^30^ and convert dietary polyphenols into bioactive metabolites with antimicrobial and anti-inflammatory properties.^31,32^ They also strengthen intestinal barrier integrity by enhancing mucus production, secreting antimicrobial peptides such as bacteriocins, thereby limiting pathogen colonization and translocation.^33^ Dysbiosis of this complex intestinal ecosystem can negatively impact overall health.^34^ Given the central role of the gut microbiome and its metabolites in maintaining intestinal barrier function and immune homeostasis, considerable interest has emerged in identifying dietary bioactive compounds capable of supporting these processes.^35^ Among these, lactoferrin has gained attention for its multifaceted effects on gut health, microbial composition, and immune regulation.^36^

Lactoferrin (LF) is an iron binding glycoprotein, primarily present in milk and various secretory fluids such as saliva, tears, and nasal secretions.^37^ It is also present in neutrophil granules and produced by hematopoietic tissue of bone marrow.^38^ It exhibits a wide range of biological activities, including supporting healthy immune function, helping to maintain balanced inflammatory responses, contributing to antioxidant status, promoting a favorable microbial balance, and supporting iron homeostasis.^36,39^ In addition, LF also influences cellular responses such as cell cycle regulation, proliferation, and cellular differentiation, thereby enhancing wound healing and barrier integrity.^40–42^

Numerous in vitro and in vivo studies have demonstrated that LF functions as both an intestinal barrier protector and intestinal inflammation regulator.^40,43,44^ Its positively charged structure enables binding to negatively charged components on immune cells and pathogens thereby triggering pathways that regulate cellular responses such as activation, differentiation, and proliferation.^45^ LF attenuates LPS-induced inflammatory responses by reducing the expression of pro-inflammatory cytokines such as tumor necrosis factor-α (TNFα), interleukin (IL)-1β, and IL-6.^46^ In parallel, it enhances intestinal barrier integrity by upregulating tight junction-associated proteins such as occludin and ZO-1.^43^ In addition, LF modulates gut microbiota composition, supports gastrointestinal development, and has been associated with increased production of SCFAs, collectively contributing to improved intestinal homeostasis.^47–49^

The widespread use of human lactoferrin (hLF) as a nutritional ingredient has been historically constrained by its limited natural availability, as commercial production from human milk, its primary source, is not scalable. Precision fermentation offers a sustainable and scalable alternative for producing recombinant human lactoferrin while overcoming supply limitations associated with human-derived protein.^50^ The recombinant protein evaluated in this study was effera^®^ hLF, developed by Helaina Inc. using an industrial scale *Komagataella phaffii* precision fermentation platform.^51^ Comprehensive biochemical and biophysical characterization has demonstrated that effera^®^ closely recapitulates the overall structural properties of native human milk lactoferrin (hmLF), including secondary and tertiary structure, globular conformation, molecular weight, thermal stability, and glycan location to the three conserved N-linked glycosylation sites.^51^ effera^®^ was developed as a food ingredient and dietary supplement engineered to closely resemble the structural and biological properties of native hLF, thereby enabling broader evaluation of its potential health benefits across diverse populations.^51,52^

To characterize the gastrointestinal activity of effera^®^, we used the validated ex vivo SIFR^®^ technology pipeline to assess SCFA production, microbial composition and host-microbiome interactions. This platform preserves donor-specific gut microbial communities and enables integrated evaluation of microbial metabolism and host-relevant functional responses. Importantly, findings generated using this platform have previously demonstrated predictive relevance for clinical changes in microbiome composition, gut barrier integrity and immune function.^53^ The present study was therefore undertaken to determine whether precision fermentation-derived effera^®^ hLF modulates microbial metabolism and intestinal barrier function through mechanisms consistent with the proposed gut-health benefits of native hmLF.

## Materials and Methods

### Test compounds

Test products evaluated were: effera^®^ (recombinant hLF) (Helaina Inc, NY, USA), bovine lactoferrin (bLF) (Lactoferrin Co, Brisbane, AUS), skim milk (Select Custom Solutions, WI, USA). Native hmLF was isolated from donor breast milk at Helaina Inc. A no-substrate control (NSC), in which the microbial inoculum was grown in absence of additional test products, yet in presence of an optimized nutritional medium, was used as a control.

### Isolation of native hmLF

Donor human milk was collected from a single donor and frozen at -20 °C prior to use. The human milk was thawed at 25 °C for 1-2 h and hmLF was isolated using the following procedure adapted from Boseman-Finkelstein and Finkelstein^54^: samples were centrifuged at 10,000 x g for 30 min, the liquid fraction (infranatant) was collected and the fat layer discarded. The resultant liquid fraction was adjusted to pH 4.7 using 1 M hydrochloride (Sigma-Aldrich, St. Louis, MO, USA) before incubation in a bead bath at 40 °C for 30 min for casein precipitation. The solution was then centrifuged at 10,000 x g for 30 min at 4 °C to separate supernatant from the precipitated casein.

The supernatant was decanted and purified via ultrafiltration/diafiltration using a 30 kDa molecular weight cut-off filter (MWCO, TangenX PRO pD Casette [Repligen, Waltham, MA, USA] on a TFF system (KrosFlo KR2i TFF [Repligen])) and then cation exchange chromatography (with a HiTrap SP HP Column (5 mL, Cytiva, Marlborough, MA, USA)) on an AKTA Pure system (Cytiva). The column was equilibrated in 50 mM sodium phosphate (pH 7.5, MilliporeSigma, Burlington, MA, USA), and the sample was applied to the column. The column was then washed with 50 mM sodium phosphate (pH 7.5) 0.1% triton x 114 for ten column volumes followed by 50 mM sodium phosphate (pH 7.5) for ten column volumes before being eluted with 50 mM sodium phosphate and 1 M sodium chloride (pH 7.5, MilliporeSigma) via an elution gradient (0 to 100% 1 M sodium chloride) for 20 column volumes. The purified product was then simultaneously buffer-exchanged and concentrated with a 30 kDa MWCO filter (15 mL of filter volume, MilliporeSigma). This procedure yielded human milk lactoferrin with a purity of greater than 95% as measured by HPLC and SDS-PAGE as described previously.^51^

### Simulation of upper GIT

Upper gastrointestinal digestion/absorption and colonic fermentation were performed as recently described.^55,56^ Products (or distilled H_2_O for NSC) were subjected to oral, gastric and small intestinal digestion according to the INFOGEST 2.0 method. To ensure compatibility with colonic incubations, modifications were implemented, such as the removal of oxygen and simulation of small intestinal absorption using a dialysis membrane with a molecular weight cut-off of 1 kDa.

### Ethics statement

Fecal samples were collected according to a procedure approved by the Ethics Committee of the University Hospital Ghent (reference number BC-09977).

### Fecal microbiota sourcing

To simulate biorelevant gut microbiota ex vivo using the SIFR^®^ technology pipeline (Cryptobiotix, Ghent, Belgium), fresh fecal samples were obtained from six human adults: test subjects 1 (male, 34), 2 (female, 49.5), 3 (male, 42.1), 4 (female, 30.4), 5 (female, 31.5), and 6 (female, 53.2). The selection criteria for the donors were as follows: 25-65 years of age, no antibiotic use 3 months prior to participation, no gastro-intestinal disorder (cancer, ulcers, inflammatory bowel disease (IBD)). Participants had not used antibiotics within three months before donation and had no history of gastrointestinal disorders (cancer, ulcers, IBD). A fat and fiber behavior questionnaire was completed by the donors. Based on this, the fat index, fiber index and total index were calculated respectively based on answers related to fat consumption, fiber consumption and both according to Reeves et al.^57^

### Ex vivo simulation of colonic fermentation via SIFR^®^ technology pipeline

Colonic simulation was performed as described recently.^53,56^ Briefly, individual bioreactors were processed in parallel in a bioreactor management device (Cryptobiotix, Ghent, Belgium). Each bioreactor contained 5 mL of nutritional medium-fecal inoculum blend supplemented with test products derived from the upper GIT simulation, then sealed individually, before being rendered anaerobic. Blend M0017 was used for preparation of the nutritional medium (Cryptobiotix, Ghent, Belgium). After preparation, bioreactors were incubated under continuous agitation (140 rpm) at 37 C (MaxQ 6000, Thermo Fisher Scientific, Merelbeke, Belgium).

Twelve study arms were tested for each test subject (n=6):

**Table 1.** Study arms, test products, respective test doses along with abbreviations that were used to refer to the study arms along the report.

| # | Study arm | Dose | Terminology |
| --- | --- | --- | --- |
| 1 | No-substrate control | N/A | NSC |
| 2 | Skimmed milk | 50 | SM_50 |
| 3 | Skimmed milk | 1000 | SM_1000 |
| 4 | Human lactoferrin | 1000 | hmLF_1000 |
| 5 | effera<sup>®</sup> | 500 | effera<sup>®</sup>_50 |
| 6 | effera<sup>®</sup> | 100 | effera<sup>®</sup>_100 |
| 7 | effera<sup>®</sup> | 300 | effera<sup>®</sup>_300 |
| 8 | effera<sup>®</sup> | 500 | effera<sup>®</sup>_500 |
| 9 | effera<sup>®</sup> | 1000 | effera<sup>®</sup>_1000 |
| 10 | Bovine lactoferrin (bLF) | 100 | bLF_100 |
| 11 | Bovine lactoferrin (bLF) | 500 | bLF_500 |
| 12 | Bovine lactoferrin (bLF) | 1000 | bLF_1000 |

At 0 h (NSC only) and after 48 h of incubation, upon gas pressure measurement in the headspace, liquid samples were collected for subsequent analysis of key fermentative parameters, microbial composition, and host-microbiome interactions.

### Key fermentative parameters

Concentrations of short-chain fatty acids (SCFA; acetate, propionate, butyrate, and valerate) and branched-chain fatty acids (BCFA) were determined via gas chromatography with flame ionization detection (Trace 1300, Thermo Fisher Scientific) upon diethyl ether extraction as previously described.^58^ pH was measured using a calibrated pH electrode (Hanna Instruments Edge HI2002, Temse, Belgium) and the accumulation of gases in the headspace was determined by a needle connected to a pressure meter as previously described.^55^

### Microbial composition analysis

Quantitative insights were obtained by correcting proportions (%; 16S rRNA gene profiling) with total counts (cells/ml; flow cytometry), resulting in estimated cells/ml of different taxa. To determine total counts, samples were diluted in anaerobic phosphate-buffered saline (PBS), followed by cell staining with SYTO 16 at a final concentration of 1 µM, and counted via the NovoCyte Quanteon flow cytometer (Agilent). Data were analyzed using NovoExpress, version 1.6.2. DNA extraction and sequencing analysis were performed as previously described.^55,59^ Briefly, DNA was extracted via the SPINeasy DNA Kit for Soil (MP Biomedicals, Eschwege, Germany), according to manufacturer’s instructions. Library preparation and sequencing were performed on an Illumina MiSeq platform (Illumina, San Diego, CA, USA) with v3 chemistry. 16S rRNA gene V3-V4 hypervariable regions were amplified using primers 341F (5’-CCTACGGGNGGCWGCAG-3’) and 785Rmod (5’-GACTACHVGGGTATCTAAKCC-3’). Preprocessing and OTU (operational taxonomic unit) picking from amplicons was performed with Mothur v1.35.155. Additionally, representative OTU sequences were annotated with the RDP-classifier using GTDB v220. If missing, a species indication was curated based on Silva v138.2.

### Gut barrier and immune modulation

Host-microbiome interaction analysis was performed to understand how the modulation of the gut microbiome (simulated using the ex vivo SIFR^®^ technology) impacted the host. A co-culture experiment with epithelial cells (human adenocarcinoma Caco-2 cell line) and immune cells (human acute monocytic leukemia THP-1 cell line, differentiated to activated macrophages by a PMA treatment) was implemented, as previously described.^56^ Briefly, upon differentiation of Caco-2 and THP-1 cells over 14 and 2 days, respectively, a co-culture model was created by covering immune cells with an epithelial layer in a permeable well insert. The impact of three types of samples on the co-culture model was evaluated: (i) four control samples including two blanks (either with or without LPS addition at 24h), and two positive controls with dexamethasone (D) or hydrocortisone (HC); (ii) direct cell exposure of unfermented test products, before digestion in the upper GIT simulation, alongside a corresponding blank control, and (iii) colonic samples obtained after 48h fermentation. These samples were centrifuged and filtered prior to being used in the co-culture assay. The host-microbiome interaction assay consisted of two phases: (i) 24 h treatment during which samples were applied on the apical side of epithelial cells to evaluate their impact on gut barrier integrity under unstressed conditions, and (ii) a subsequent additional 6h incubation in the presence of 500 ng/mL LPS to evaluate effects on barrier integrity under stressed conditions, along with potential immunomodulatory effects (at 30h). Transepithelial electrical resistance (TEER) is a widely accepted method to measure gut barrier integrity^60^ and was measured before administration of colonic samples (0h), at 24h (before LPS addition) and at 30h (6h after LPS addition). For each well, the TEER values at 24h and 30h were normalized against the value measured at 0h to correct for small differences between the wells. Immune effects at 30h were studied via cytokine/chemokine production using a Multiplex Luminex® Assay kit on the MAGPix® analyzer (IL-6, CXCL-10, IL-10, IL-1β, TNF-α) or ELISA (IL-8). In addition, expression of tight junction targets ZO-1 and occludin was measured by RT-qPCR on the cell fraction.

### Data analysis

Statistical data analysis was performed using GraphPad Prism (Version 10.4.2, GraphPad software. San Diego, CA, USA) and R (version 4.5.1; www.r-project.org) as appropriate. For the statistical evaluation of the treatment effects on key fermentative parameters, cell counts, microbial diversity (4 indices), microbial composition (phylum level), gut barrier integrity and immune markers, a linear mixed-effects model was implemented for each parameter, across six different donors. This approach accounts for both fixed effects (treatment) and random effects (individual variability). Subsequently, a *post hoc* pairwise comparison with Benjamini-Hochberg corrections of p-values was applied to identify significant differences between the effects of the treatments. Statistical differences vs different references from the *post hoc* tests were visualized on the violin plots. A treatment response was considered consistent when the taxon was detected in at least four donors and showed the same direction of change relative to the NSC in every donor in which it was present.

## Results

### Microbiota of study subjects captured inter-donor differences in gut microbiota composition

A principal component analysis at the genus level demonstrated that there were marked differences in microbial composition between the six donors at baseline (Figure 1). The inter-donor differences in fecal microbiome composition at baseline were in line with the stratification of human gut microbiota according to the concept of unique enterotypes. Donor 2 was distinguished from the other donors by a high relative abundance of Prevotella, a defining feature of the Prevotella enterotype. Donors 5 and 6 exhibited elevated levels of *Bacteroides/Phocaeicola*, characteristic of the Bacteroides enterotype. In contrast, donor 1 showed higher abundances of *Bacillota_A* genera, particularly members of the *Lachnospiraceae* and *Ruminococcacea*e families, consistent with the *Ruminococcus* enterotype. Finally, donor 4 displayed a notably higher abundance of Bifidobacterium compared to the other donors. These initial findings show that the fecal microbiota of the six healthy adults encompassed relevant phenotypic differences and the broad spectrum of in vivo microbiota composition, thus capturing representative variation across a population.

**Figure 1:**
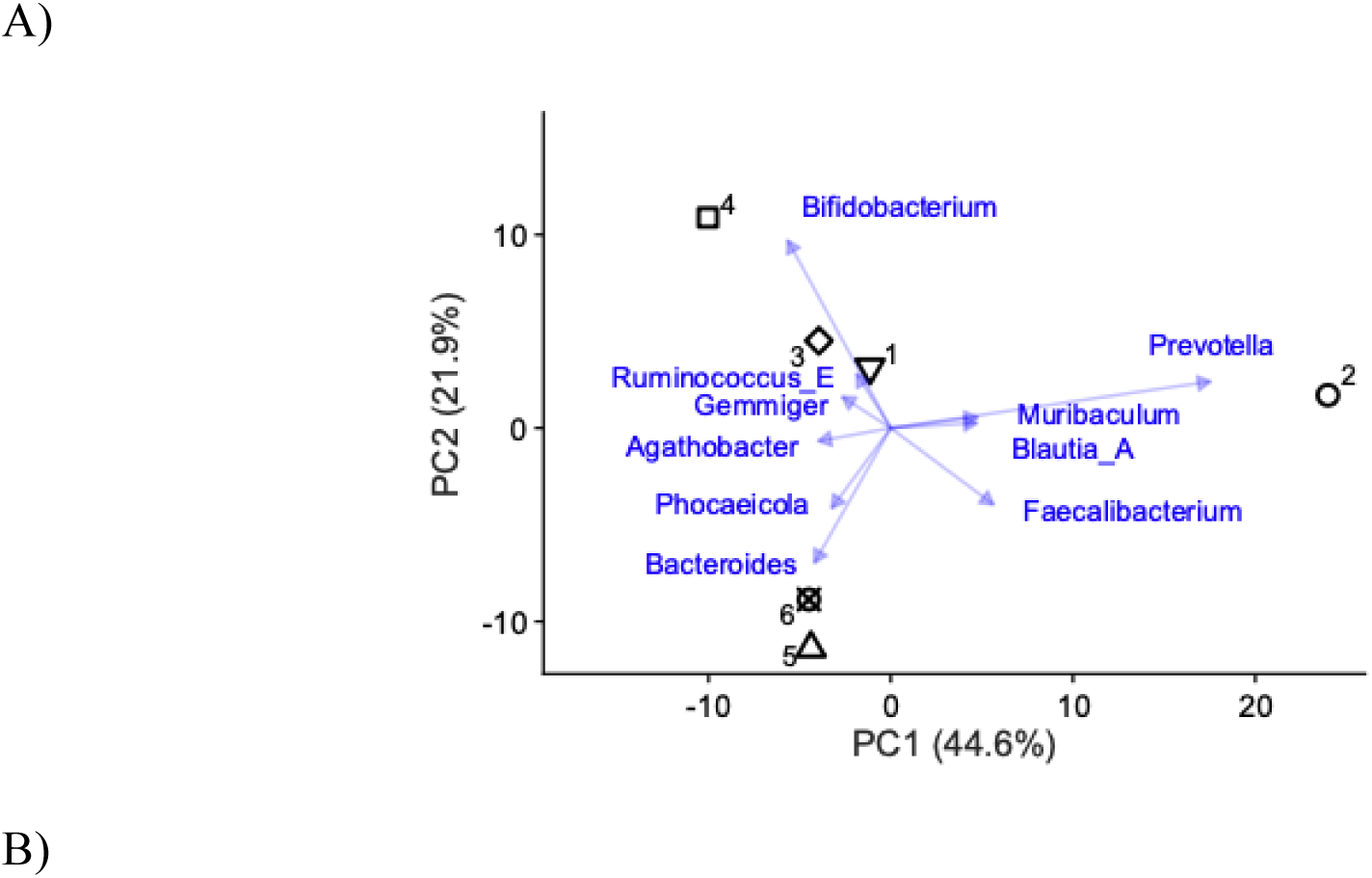
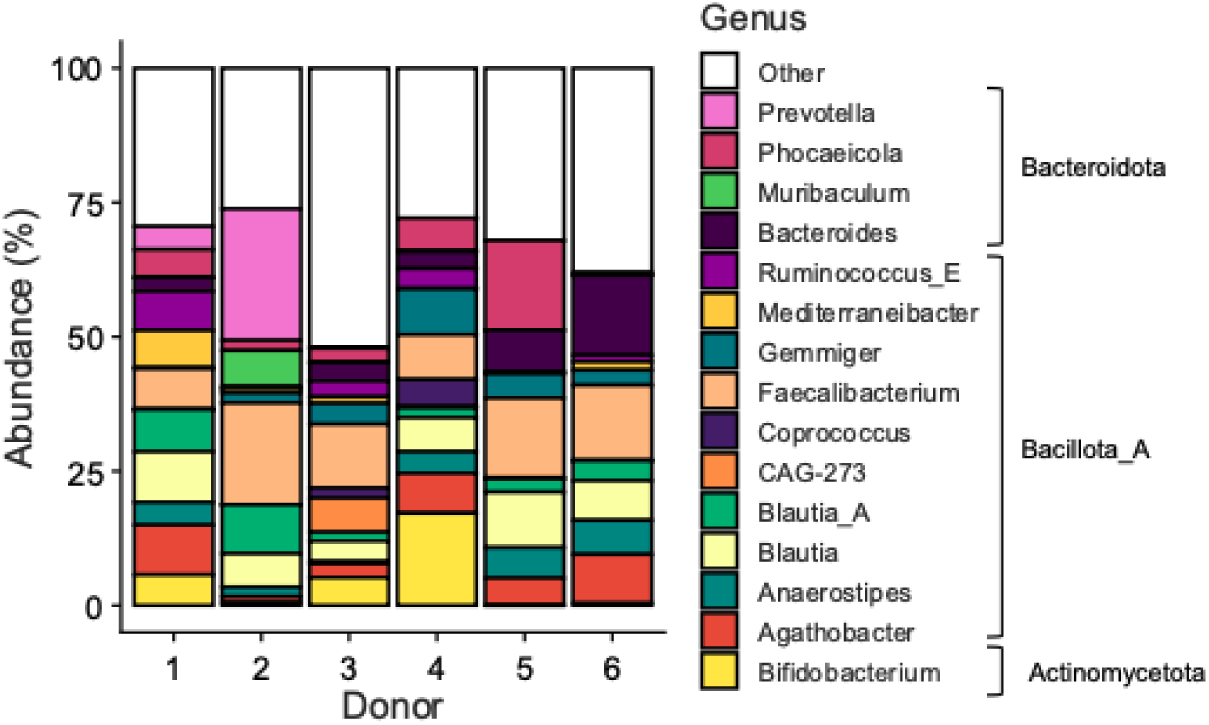
(A) Principal component analysis (PCA) summarizing the microbial community composition of the six human adults that provided a fecal donation for the current study. The PCA was calculated based on the (centered) abundances (%) of the microbial genera, as quantified via 16S rRNA gene profiling. (B) Abundances (%) of the key genera of the different fecal microbiota.

### effera^®^ modulates microbial fermentation and metabolic output

Microbial fermentation activity was assessed by measuring changes in pH, gas production, SCFAs, and branched-chain fatty acids (BCFAs) across increasing concentrations of effera^®^. effera^®^ significantly altered multiple fermentation-derived metabolic outputs in a dose-dependent manner. Total SCFA production was significantly increased compared to NSC starting at 50 mg/day, with early effects primarily driven by increases in butyrate and propionate (Figure 2 A-C). At doses ≥300 mg/day, significant increases in valerate (Figure 2D) and at ≥500 mg/day a significant increase in acetate were also observed, with the strongest overall SCFA responses observed at 500 mg/day and 1000 mg/day (Figure 2E). BCFA production was also significantly elevated across all effera^®^ doses, with more pronounced effects emerging from 300 mg/day onward (Figure 2F). effera^®^ significantly increased gas production in a dose-dependent manner, with effects first observed at 100 mg/day and progressively increasing at higher doses (Figure 2G). This increase in the gas production was consistent with enhanced overall microbial fermentative activity and paralleled the observed increases in SCFA and BCFA production. effera^®^ did not induce significant changes in the pH at any tested dose compared with the NSC (Figure 2 H) Bovine LF was evaluated in parallel as a comparator and similarly increased SCFA and BCFA production across tested doses (Figure 2 A-F). Significant increase in the gas production was observed in a dose-dependent manner without affecting pH (Figure 2 G-H). Skim milk and native hmLF at 1000 mg/day significantly increased total SCFA, along with significant increases in acetate, butyrate, propionate, and valerate, along with an increase in the gas production with no changes in the pH. No significant changes were observed at 50 mg/day of skim milk. (Figure 2 A-H)

**Figure 2:**
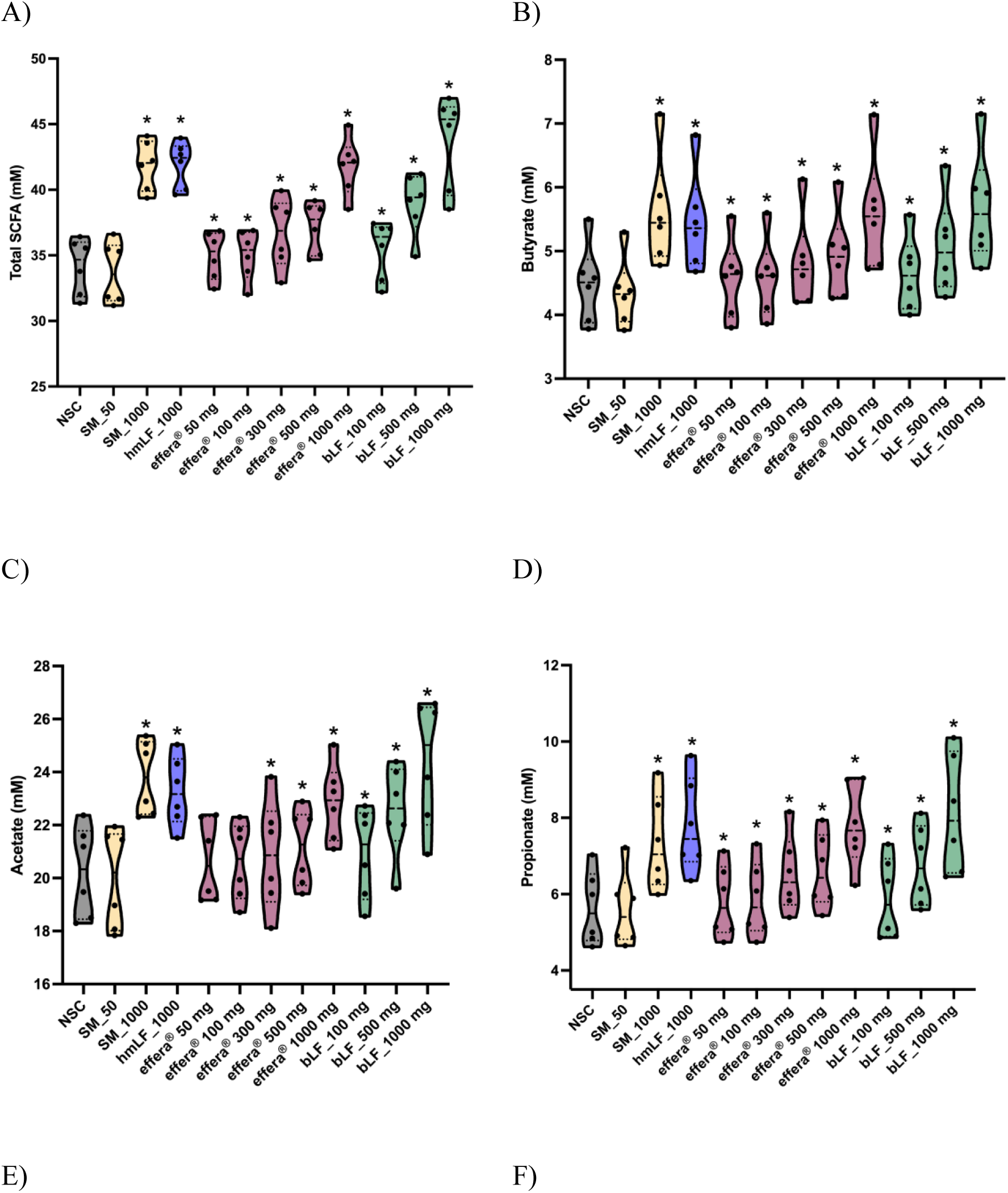
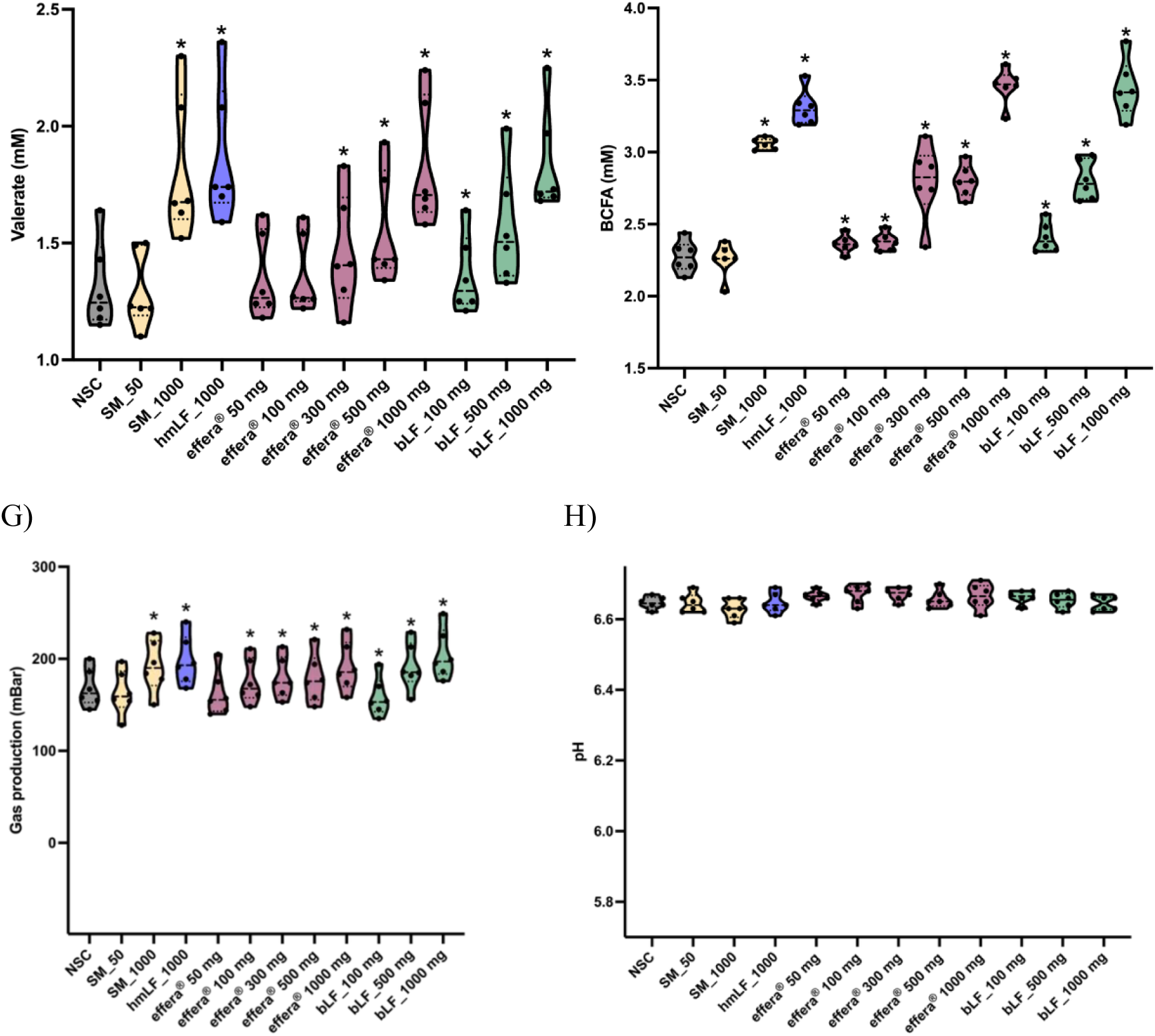
The impact of the test products on (A) total SCFA, (B) butyrate, (C) acetate, (D) propionate, (E) valerate, (F) BCFA, (G) gas production, (H) pH for healthy at 48h. Statistical differences between NSC and the individual treatments. Violin plots represent responses generated using microbiota from six healthy adult donors. Statistical analysis was performed using linear mixed-effects model with donor included as a random effect, followed by *post-hoc* comparisons with Benjamini-Hochberg adjustment for multiple testing. Adjusted P<0.05 was considered statistically significant. * indicates adjusted P<0.05 versus the non-supplemented control.

### effera^®^ increased gut microbial cell density while maintaining diversity

To assess the broader impact of effera^®^ on gut microbial complexes, ex vivo colonic fermentation was conducted using fecal samples from six adult donors for 48 h. Bacterial cell density was quantified using flow cytometry and microbial growth was analyzed across treatment groups. effera^®^ significantly increased the bacterial cell density in a dose-dependent manner starting at 100 mg/day (Figure 3). Bovine LF significantly increased the bacterial cell density starting at 500 mg/day (Figure 3). The increase in gut microbiome cell density was significantly higher than bLF at all the doses tested for effera^®^. effera^®^ induced an increase of greater magnitude and earlier onset in gut microbial cell density compared to bLF. Both native hmLF and skim milk at 1000 mg/day significantly increased the bacterial cell density (Figure 3). While the treatments significantly increased the bacterial cell density, they did not alter the diversity or evenness of the microbial community.

**Figure 3:**
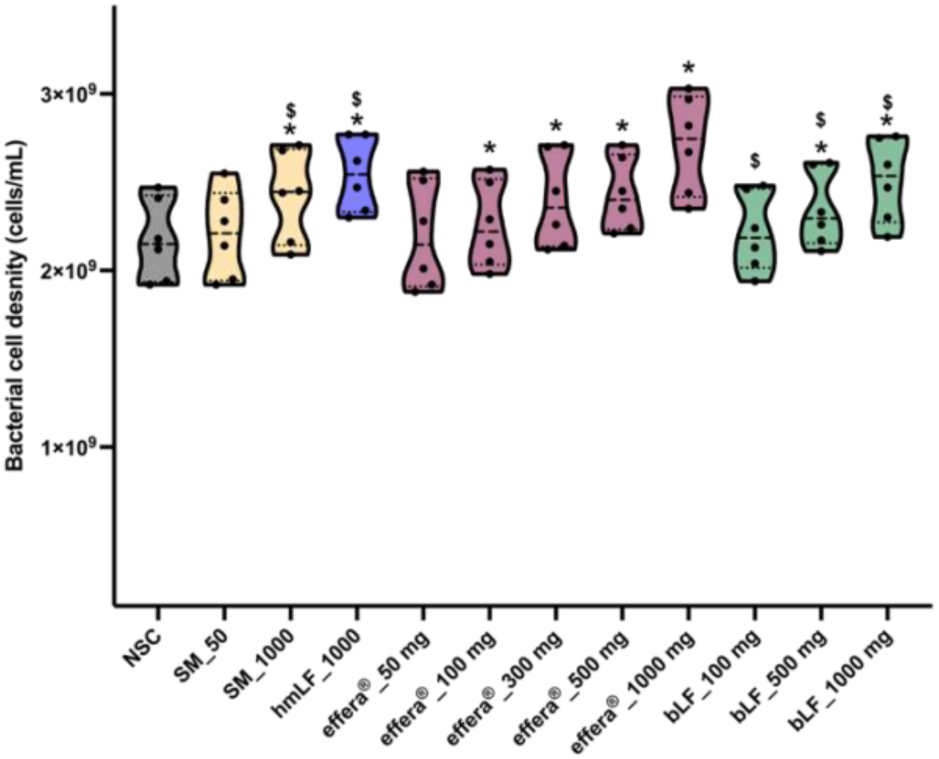
The impact on total bacterial cell counts of the test products compared to a non-supplemented control (NSC). Violin plots represent responses generated using microbiota from six healthy adult donors. Statistical analysis was performed using linear mixed-effects model with donor included as a random effect, followed by *post-hoc* comparisons with Benjamini-Hochberg adjustment for multiple testing. Adjusted P<0.05 was considered statistically significant. * indicates adjusted P<0.05 versus the non-supplemented control.

To determine whether these increases in microbial biomass were accompanied by microbial shifts, three distinct alpha diversity metrics were assessed using Chao1, Shannon, and reciprocal Simpson diversity index. The Chao1 diversity index is a measure of species richness, estimating the total number of species present within a sample, and the Shannon diversity index accounts for both species’ richness and evenness. No significant differences were observed in Chao1 species richness or the Shannon diversity index across treatments or doses, indicating that overall microbial diversity and evenness remained stable (data not shown). These results suggest that effera^®^ increased microbial biomass without reducing alpha diversity.

### effera^®^ induces phylum and family level shifts in microbial composition

The impact of effera^®^ on gut microbial community composition was evaluated at higher taxonomic resolutions, including phylum and family levels. At the phylum level, effera^®^ induced distinct and dose-dependent shifts compared to NSC characterized by a consistent increase in both *Bacillota_A* and *Bacteroidota*. (Figure 4A-B). Significant increases in the abundance of *Bacillota_A* phylum and *Bacteroidota* were observed with effera^®^ treatment starting at 300 mg/day. Bovine LF also significantly increased the *Bacillota_A* at 500 mg/day dose (Figure 4B). However, the magnitude of increase was lower than was observed with effera^®^ at all the tested doses. In contrast, there were no statistically significant differences in *Bacteroidota* observed with bLF even at the highest dose tested (Figure 4A). Native hmLF significantly increased both the phyla while skim milk had no effect on them (Figure 4A-B).

**Figure 4:**
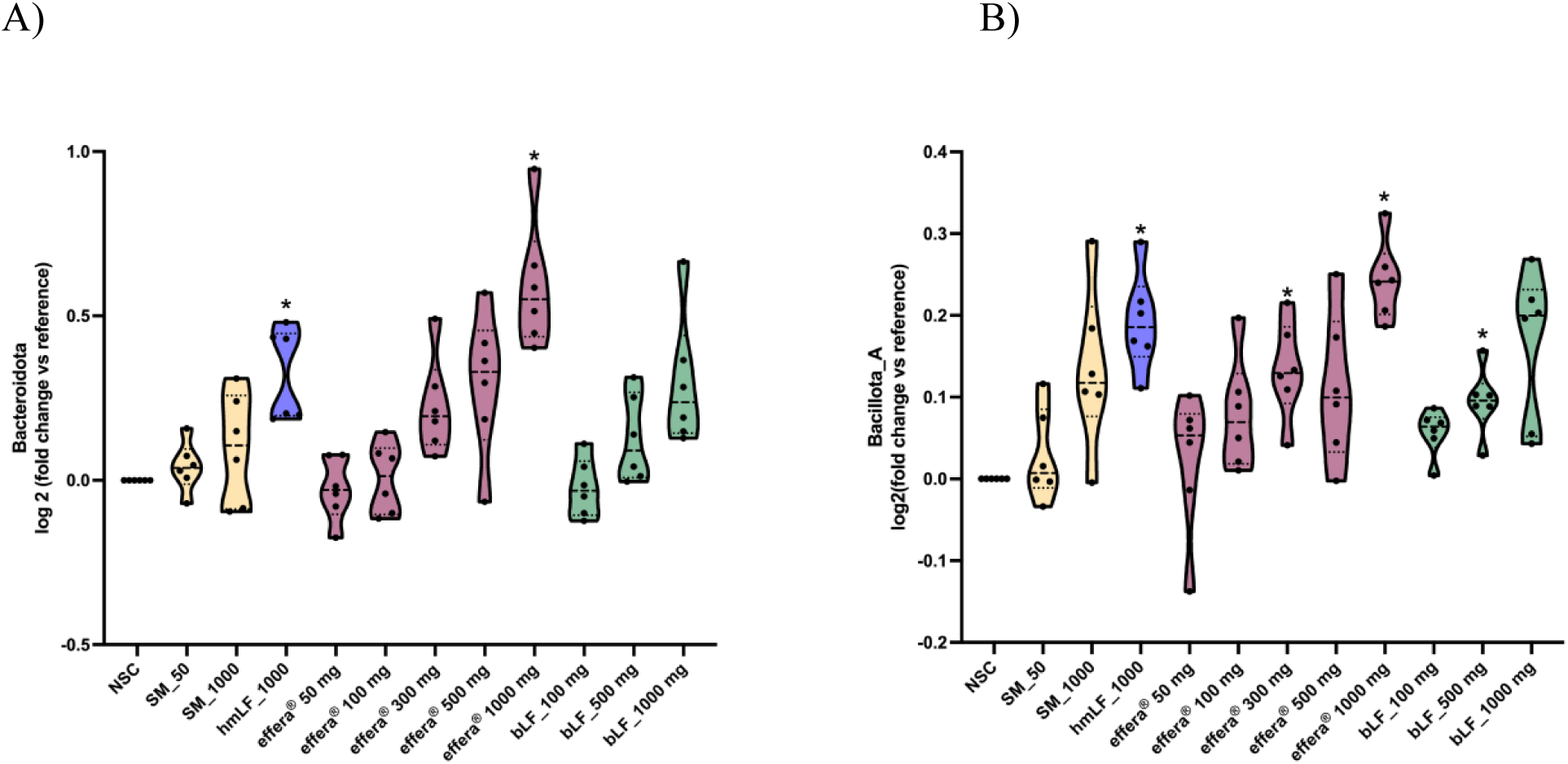
The impact of the test products on bacterial phyla compared to NSC. Violin plots represent responses generated using microbiota from six healthy adult donors. Statistical analysis was performed using linear mixed-effects model with donor included as a random effect, followed by *post-hoc* comparisons with Benjamini-Hochberg adjustment for multiple testing. Adjusted P<0.05 was considered statistically significant. * indicates adjusted P<0.05 versus the non-supplemented control.

At the family level, the increase in *Bacillota_A* was primarily due to the enrichment of Oscillospiraceae. Within the *Bacteroidota* phylum, increased abundance was driven by enhancement of families *Bacteroidaceae*, *Barnesiellaceae*, *Rikenellaceae*, and *Tannerellaceae* indicating a broad activation of *Bacteroidot*a-associated families represented as a heat map (Figure 5).

**Figure 5:**
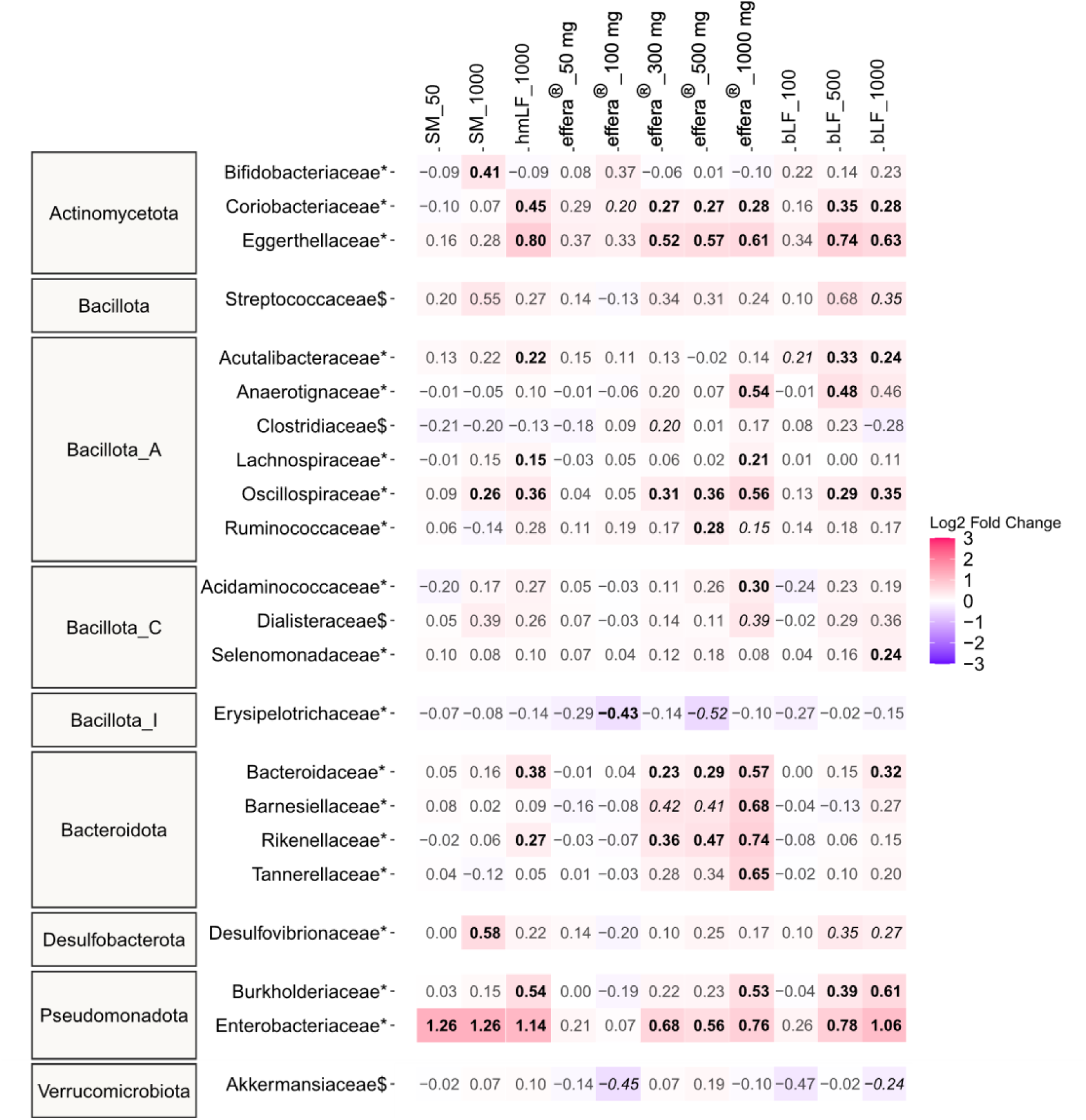
Effects of the test products on bacterial families after 48 h of colonic incubation. The heatmap includes families that were significantly affected (p_adjusted_ < 0.05; *) or showed a consistent treatment response ($) for at least one treatment. Data are presented as mean log_2_ fold changes in absolute abundance relative to the NSC (log_2_[treatment/NSC]) across six donors. Values are shown in bold when the treatment effect was statistically significant and in italics when a consistent increase or decrease was observed. Together, these results demonstrate that effera^®^ selectively modulates key dominant microbial lineages at both the phylum and family levels in a dose-dependent manner, without disrupting global community structure. To further understand these compositional changes at higher taxonomic resolution, species level analyses were performed and represented as a heat map (Figure 6).

**Figure 6:**
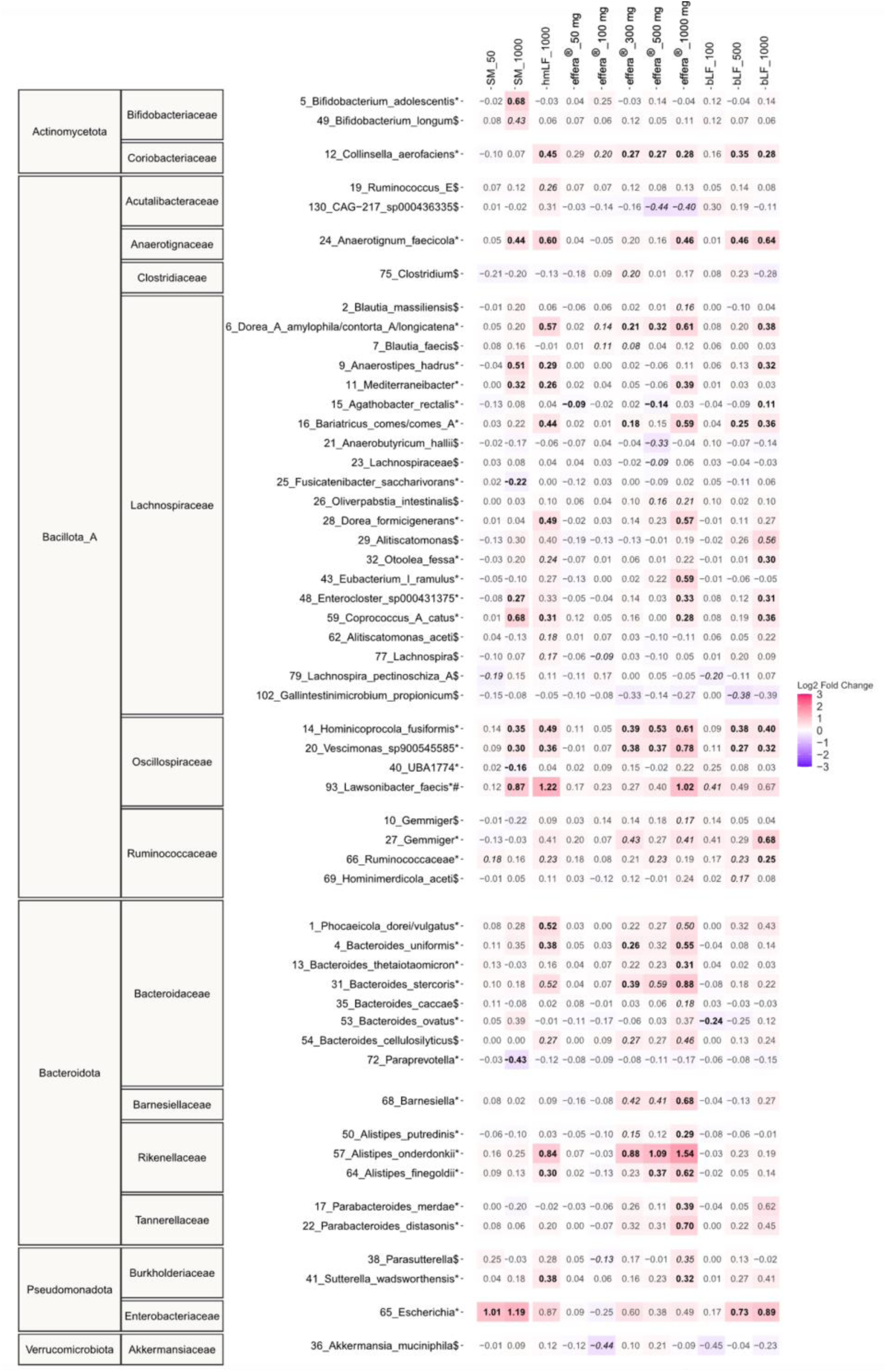
Effects of the test products on OTUs assigned to specific microbial species after 48 h of colonic incubation. The heatmap includes OTUs that were significantly affected (p_adjusted_ < 0.05; *) or showed a consistent treatment response ($) for at least one treatment. Data are presented as mean log_2_ fold changes in absolute abundance relative to the NSC (log_2_[treatment/NSC]) across six donors. Values are shown in bold when the treatment effect was statistically significant and in italics when a consistent increase or decrease was observed.

In the *Oscillospiraceae* family, effera^®^ treatment resulted in distinct increases in *Hominicoprocola_fusiformis*, *Lawsonibacter_faecis* and *Vescimonas_sp900545585* species. Among the Barnesiellaceae family, an increase in *Barnesiella* species was observed. Within the Bacteroidaceae family, effera^®^ increased the abundance of several species including *Phocaeicola_dorei/vulgatus*, *Bacteroides_uniformis*, *Bacteroides_thetaiotaomicron*, *Bacteroides_stercoris*, and *Bacteroides_cellulosilyticus*. Additional increases were observed within the Rikenellaceae family, including *Alistipes_putredinis*, *Alistipes_onderdonkii*, and *Alistipes_finegoldii*, as well as with in Tannerellaceae family, including Parabacteroides_merdae and *Parabacteroides_distasonis*. In comparison, bLF treatment resulted in distinct increases in a limited subset of species, primarily in *Lawsonibacter_faecis*, *Barnesiella*, *Parabacteroides_merdae* and *Parabacteroides_distasonis*. Overall, effera^®^ resulted in a broader species level shift compared to bovine lactoferrin with the most pronounced increase observed at 1000 mg/day. Additional compositional changes at the family level were evaluated to further characterize the effects of effera^®^ and bLF on the gut microbiota. Treatment-related effects were observed in families such as Coriobacteriaceae, Eggerthellaceae, Anaerotignaceae, Lachnospiraceae, Acidaminococcaceae, and Burkholderiaceae. Skim milk significantly increased Bifidobacteria at 1000 mg/day, with no significant effects observed at the lower dose tested, 50 mg/day.

### effera^®^ promoted gut barrier integrity under basal (unstressed) and inflamed (stressed) conditions

The effects of effera^®^ and bLF on intestinal barrier integrity were evaluated using a Caco-2/THP-1 co-culture model. To distinguish direct product effects from those emerging during microbial fermentation, sample types were evaluated separately: intact test products before digestion and cell-free samples collected after ex vivo colonic fermentation using the SIFR^®^ technology pipeline.

Under basal (unstressed) conditions, exposure to intact products for 24 hours did not result in significant changes in transepithelial electrical resistance (TEER), indicating no direct effect of intact protein on epithelial barrier integrity (data not shown). In contrast, treatment with colonic fermentation-derived metabolites resulted in a marked enhancement of barrier function, reflected by increased TEER values. effera^®^ exhibited a clear dose-dependent response, with a significant increase in the TEER observed at the doses 300 mg/day and above compared to NSC (Figure 7A). At the highest dose tested (1000 mg/day), effera^®^ fermentation products induced a significantly greater increase in TEER compared with bLF at the at 1000 mg/day dose, indicating a stronger barrier-enhancing effect under basal conditions. Fermentation-derived metabolites of native hmLF significantly increased TEER compared to NSC at 1000 mg/day under basal conditions (Figure 7A). Fermentation-derived metabolites of skim milk increased TEER at 1000 mg/day, with no effect observed at 50 mg/day.

**Figure 7:**
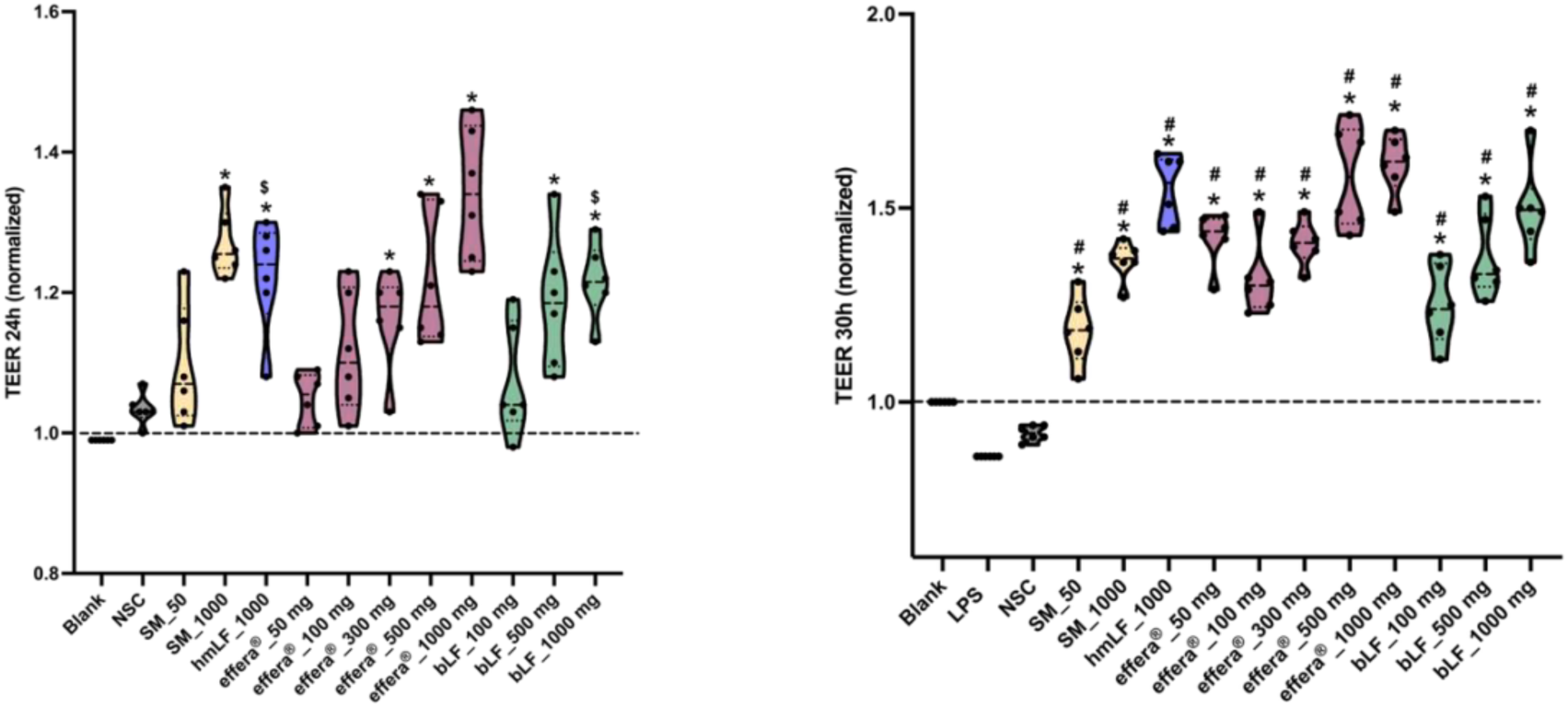
Effects of post-colonic fermentation samples on intestinal epithelial barrier integrity under basal and inflammatory conditions. Test products were subjected to upper gastrointestinal digestion followed by 48 h of ex vivo colonic fermentation. Cell-free fermentation supernatants were subsequently applied to a Caco-2/THP1co-culture model, and TEER values were assesed under A) basal conditions B) LPS-induced inflammatory conditions. Violin plots represent responses generated using microbiota from six healthy adult donors. Statistical analysis was performed using linear mixed-effects model with donor included as a random effect, followed by *post-hoc* comparisons with Benjamini-Hochberg adjustment for multiple testing. Adjusted P<0.05 was considered statistically significant. * indicates adjusted P<0.05 versus the NSC and, # indicates adjusted P<0.05 versus LPS alone, and $ indicates adjusted P<0.05 versus effera^®^ at respective dose.

Barrier function was further evaluated under inflammatory conditions induced by LPS challenge. LPS exposure significantly reduced TEER, confirming disruption of epithelial barrier integrity (Figure 7B). While all samples were assessed for their ability to mitigate LPS-induced barrier dysfunction, fermentation-derived metabolites of test products again displayed the most pronounced barrier enhancing effects. effera^®^ significantly restored TEER at doses as low as 50 mg/day and demonstrated an increase in TEER across the tested range (Figure 7B). Under the LPS-challenged condition, skim milk displayed a modest effect at both doses but significantly underperformed compared to the barrier enhancement driven by effera^®^. Native hmLF at 1000 mg/day significantly restored epithelial barrier integrity, with TEER values comparable to 1000 mg/day of effera^®^ demonstrating similar barrier enhancing effects under LPS-challenged conditions (Figure 7B). Collectively, these findings demonstrate that microbial-derived fermentation metabolites of effera^®^ increase epithelial barrier integrity under both basal and LPS-challenged conditions, with enhanced barrier support under stress, and improved functional activity compared to bLF in this model system.

### effera^®^ fermentation-derived products enhance tight junction-associated gene expression

To further characterize the effects of effera^®^ on epithelial barrier function, the expression of the tight junction-associated genes TJP1, encoding ZO-1, and OCLN, encoding occludin, was evaluated following LPS challenge. LPS significantly reduced TJP1 mRNA expression compared with NSC. Treatment with fermentation-derived products of effera^®^ significantly increased TJP1 expression from 50 mg/day onward compared with LPS alone, with the largest response observed at 1000 mg/day (Figure 8A). Bovine LF, native hmLF, and skim milk also increased TJP1 expression under several treatment conditions. The consistent response across the effera^®^ dose range supports a robust effect on a key tight junction-associated pathway following inflammatory challenge.

**Figure 8:**
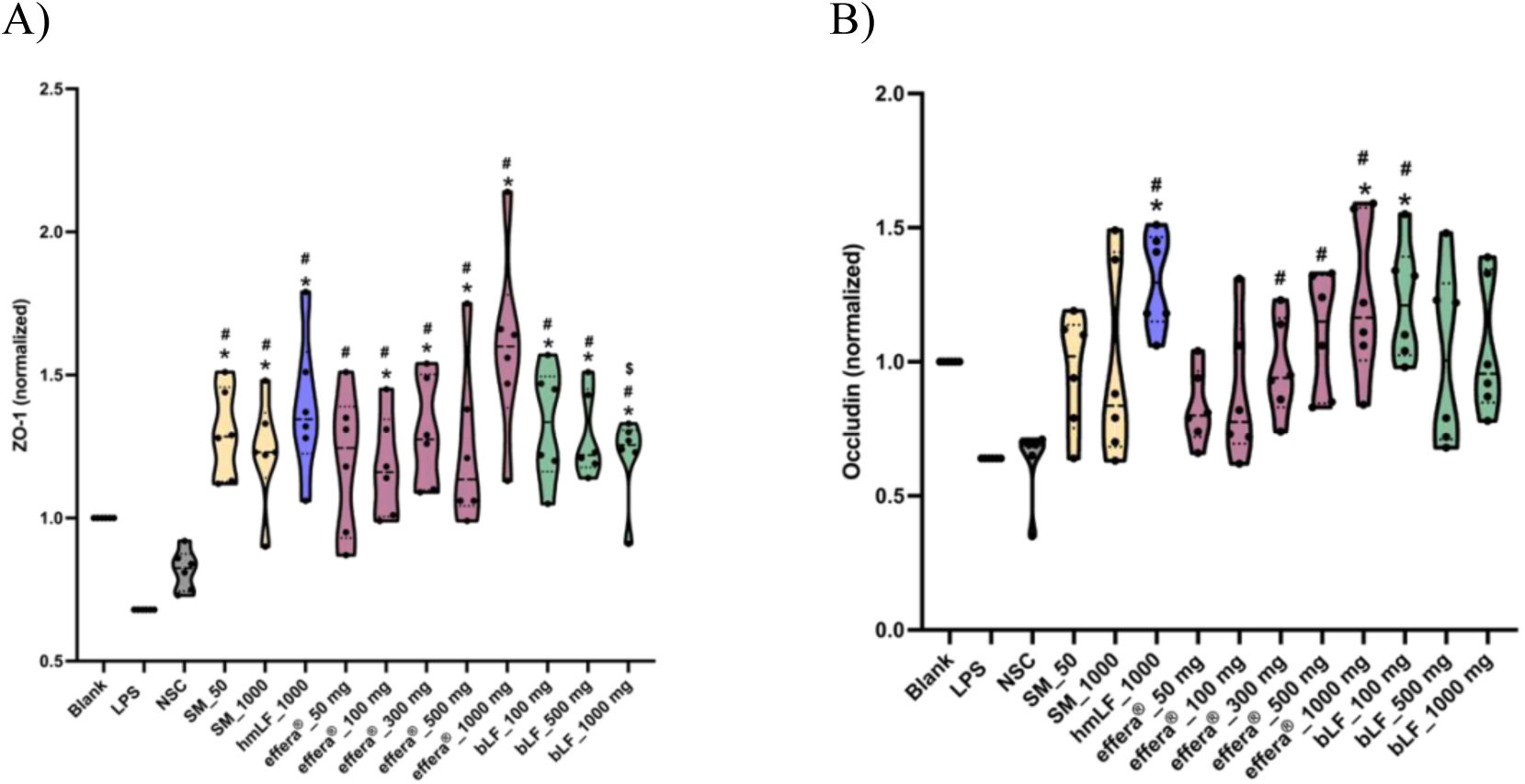
Effects of post-colonic fermentation samples on tight junction-associated gene expression following LPS challenge. Following 24 hours of exposure to filtered fermentation supernatants, the Caco-2/THP-1 co-culture model was challenged with LPS for 6 hours. Relative mRNA expression of A) TJP1, encoding ZO-1, and B) OCLN, encoding occludin was measured by RT-qPCR. Violin plots represent responses generated using microbiota from six healthy adult donors. Statistical analysis was performed using linear mixed-effects model with donor included as a random effect, followed by *post-hoc* comparisons with Benjamini-Hochberg adjustment for multiple testing. Adjusted P<0.05 was considered statistically significant. * indicates adjusted P<0.05 versus the NSC, and, # indicates adjusted P<0.05 versus LPS alone, and $ indicates adjusted P<0.05 versus effera^®^ at respective dose.

Fermentation-derived products of effera^®^ also significantly increased OCLN mRNA expression from 300 mg/day onward compared with LPS alone (Figure 8B). Native hmLF significantly increased OCLN mRNA levels at the tested dose (1000 mg/day). In contrast, bLF significantly increased OCLN expression only at 100 mg/day, with no significant effects at higher doses tested, while skim milk did not significantly alter OCLN expression. Together, these findings show that effera^®^’s fermentation-derived products enhance the transcription of key tight-junction-associated genes under inflammatory conditions. The coordinated increases in TJP1 and OCLN expression complement the observed improvement in TEER and support a microbiome-mediated barrier-enhancing effect of effera^®^.

### effera^®^ fermentation-derived products attenuate LPS-induced chemokine responses

Immunomodulatory effects of effera^®^ were evaluated following a 6-hour LPS challenge. The secretion of key cytokines and chemokines, including TNFα, IL1β, IL-6, IL-8, IL-10, and CXCL-10 was assessed. Among the biomarkers evaluated, CXCL-10 demonstrated the most pronounced response to the treatment. Fermentation-derived products of effera^®^ significantly reduced LPS-induced CXCL10 secretion across the tested dose range (Figure 9A). These findings indicate attenuation of chemokine signaling involved in inflammatory cell recruitment and activation. A comparable response in the decrease of IL-8 levels was observed among all treatment groups following LPS challenge (Figure 9B). Together, these results demonstrate that fermentation-derived products of effera^®^ selectively modulate LPS-induced chemokine responses, complementing the observed improvements in epithelial barrier integrity.

**Figure 9:**
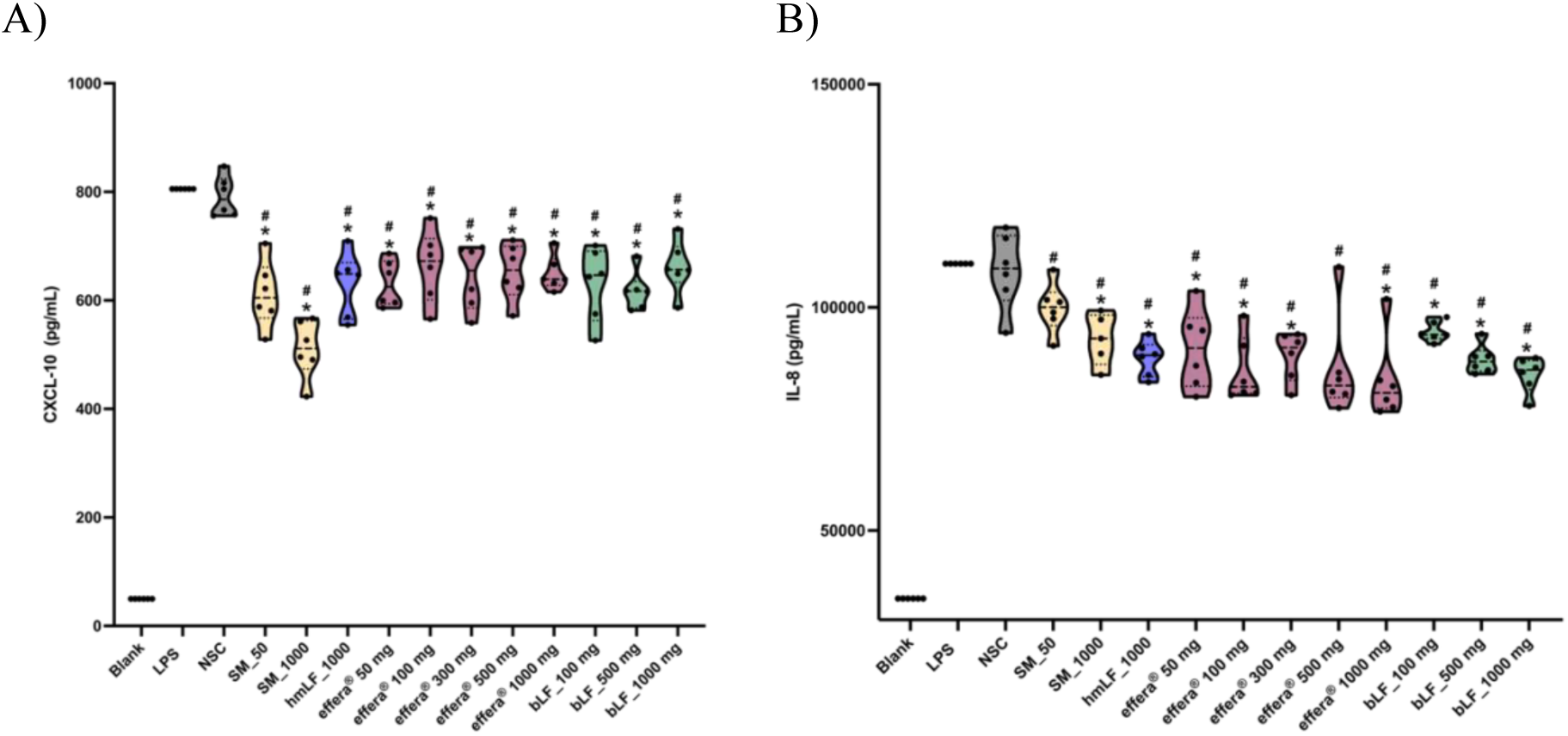
Effects of post-colonic fermentation samples on LPS-induced chemokine secretion in the Caco-2/THP-1 co-culture model. Following 24 hours of exposure to filtered fermentation supernatants, cells were challenged with LPS for 6 hours. Concentrations of A) CXCL10 and B) IL-8 levels were measured in the culture supernatant. Violin plots represent responses generated using microbiota from six healthy adult donors. Statistical analysis was performed using linear mixed-effects model with donor included as a random effect, followed by *post-hoc* comparisons with Benjamini-Hochberg adjustment for multiple testing. Adjusted P<0.05 was considered statistically significant. * indicates adjusted P<0.05 versus the NSC, and, # indicates adjusted P<0.05 versus LPS alone.

## Discussion

The present study reports that effera^®^, a recombinant hLF produced through precision fermentation, exerts broad effects on gut microbial activity, gut barrier integrity and immune response. Skim milk, bLF, and native hmLF were used as comparators. Skim milk was included as a matrix comparator to distinguish LF-specific effects from the dairy matrix. Bovine LF represented the incumbent commercial LF comparator, and native hmLF served as a physiological reference. effera^®^ promoted microbial fermentation, increased the production of beneficial metabolites such as SCFA and BCFA, and induced microbial composition shifts within key bacterial taxonomic groups associated with gut health. These microbiome-associated changes were translated to functional effects as observed with an in vitro epithelial/immune co-culture setup. Microbial fermented products of effera^®^ significantly enhanced TEER in both basal (unstressed) and inflammatory (stressed) conditions.

Importantly, the findings of the current study report that effera^®^ exerts its beneficial effects on intestinal health primarily through microbiome-mediated mechanisms rather than through the direct interactions of the intact protein with the intestinal epithelium. effera^®^ as an intact protein did not significantly improve epithelial barrier integrity; the most pronounced and significant effects were observed following colonic fermentation in the SIFR^®^ platform. A complementary process may be relevant during early life, when lactoferrin undergoes gastrointestinal digestion and releases potentially bioactive peptide products. Peptidomic analysis of human milk collected before and after gastric digestion in term infants identified significantly greater numbers of lactoferrin-derived peptides in the gastric samples, demonstrating that lactoferrin is actively transformed in the infant stomach.^61^ These findings complement the present study by supporting a sequential model in which gastrointestinal digestion first generates lactoferrin-derived products, followed by further microbial biotransformation into products with epithelial activity. An infant-adapted digestion and fermentation study would be valuable to determine whether similar bioactive products are generated in the developing gastrointestinal tract. It should therefore be noted that the studies evaluating LF’s effect solely in epithelial monocultures may underestimate LF’s potential in gastrointestinal systems where extensive interactions occur between dietary proteins, the gut microbiota, and the host. Fermentation-derived metabolites significantly enhanced TEER under both basal and inflammatory conditions. Bovine LF also followed a similar microbiome-dependent pattern of activity; however, the response of effera^®^ is greater than the effect of bovine LF on epithelial barrier integrity.

Under the conditions tested, native hmLF improved epithelial barrier function, serving as an appropriate human-derived LF comparator. When challenged with LPS, effera^®^ and hmLF demonstrated comparable effects in protecting the epithelial barrier integrity, demonstrating that effera^®^ maintains the functional property of hmLF under LPS-challenged conditions.

Disruption of intestinal barrier integrity is a central feature of numerous gastrointestinal disorders and also an important contributor to chronic inflammation and susceptibility to infections.^34,62,63^ The intestinal epithelium is the first line of defense between the luminal environment and the systemic circulation.^1^ It regulates selective absorption of nutrients while preventing harmful bacteria, endotoxins and immunogenic antigens.^2,3^ When the barrier gets compromised, microbes and their metabolites can penetrate the barrier and activate the immune system, resulting in continuous inflammation and further compromising the intestinal epithelium.^64^ Preservation of the intestinal barrier is critical for preventing entry of harmful bacteria and immunogenic pathogens.^64^ LPS is a structural component of gram-negative bacteria and is one of the most extensively studied initiators of inflammation. Binding of LPS to TLR4 results in activation of a proinflammatory signaling cascade and release of cytokines and chemokines.^65^ In our study, LPS challenge significantly impaired epithelial barrier integrity as demonstrated by a reduction in the TEER value and decreased the expression of tight junction-associated genes. Remarkably, post-colonic fermentation metabolites of effera^®^ effectively counteracted this effect. This finding suggests that microbiome-derived metabolites generated from effera^®^ help maintain epithelial function during inflammatory insult. effera^®^ also significantly decreased the chemokine CXCL-10 and cytokine IL-8 levels, indicating that preservation of barrier integrity was accompanied by modulation of immune response. These findings are consistent with early-life preclinical evidence supporting a role of lactoferrin in intestinal maturation and protection. Dietary bovine lactoferrin increased intestinal crypt-cell proliferation and supported intestinal growth in neonatal piglets, demonstrating biological activity within the developing gastrointestinal tract.^40^ In addition, lactoferrin attenuated LPS-induced intestinal immune-barrier damage in young mice through modulation of ELAVL1-related immune signaling pathways.^44^ Although these studies used different models and endpoints and reduced inflammatory chemokine responses observed in the present study. Together, the evidence supports the potential relevance of lactoferrin-mediated epithelial and immune protection during early life and provides a rationale for confirmation in a dedicated infant-relevant model. Although the TEER enhancement effects were more pronounced with the LPS-inflammatory challenge, improvement of epithelial barrier function was observed in basal conditions following exposure to fermentation-derived metabolites. While the basal model does not represent overt inflammation, it represents the physiological state in which the epithelium is continuously exposed to microbial metabolites, environmental stressors, food and dietary antigens. Factors such as physiological stress, sleep disturbances, change in the diet, and aging have been associated with changes in intestinal permeability, and barrier function in healthy individuals.^66–68^ Therefore strengthening the epithelial barrier under homeostatic conditions may improve the resilience of the gut. Interestingly, skim milk also demonstrated significant effect on epithelial barrier function by increasing the TEER value at 1000 mg/day under basal conditions following microbial fermentation. This response is consistent with the ability of milk derived nutrients such as lactose and casein generating components that can support the epithelial function. However, unlike effera^®^, skim milk showed limited effect under LPS-induced inflammatory condition, suggesting that increased microbial fermentation alone is not sufficient to fully protect the intestinal barrier during inflammatory stress. The enhanced barrier effects of effera^®^ under inflammatory condition may indicate that recombinant lactoferrin promotes a more targeted microbiome-host interaction capable of enhancing the intestinal epithelium resilience beyond the effects of a general dairy matrix.

The improved barrier function observed following treatment with the fermentation metabolites of effera^®^ was accompanied by significant increases in the production of SCFA. Acetate, propionate, and butyrate are the predominant SCFAs produced by the gut microbiota and play an essential role in maintaining intestinal homeostasis.^14^ Among these, butyrate is particularly well established as a regulator of intestinal barrier integrity.^69^ Butyrate serves as the primary energy source for colonocytes and has been shown to significantly upregulate tight junction proteins by increasing the expression of ZO-1 and occludin, and reducing epithelial permeability.^19,70^ Several in vivo studies have demonstrated that butyrate increases colonocyte proliferation and intestinal villus depth, and protects against epithelial injury, promoting tissue repair.^71–75^ In addition to barrier supportive effects, SCFAs have well-documented immunomodulatory properties. Butyrate has been reported to decrease pro-inflammatory cytokines and promote anti-inflammatory cytokines.^76^ In the Caco-2 cell model, propionate significantly increased TEER, with decreased permeability to macromolecules, indicating improved barrier integrity.^77^ Propionate and acetate have also been reported to have barrier protective effects against LPS-induced acute inflammation, in an in vitro co-culture study.^78^ Valerate, although less extensively characterized, has been associated with the promotion of tight barrier integrity and reducing paracellular permeability in Caco-2 cells.^79^ In addition to promoting intestinal health, SCFA’s also aid in liver lipogenesis, improving insulin sensitivity, appetite regulation, wound healing and counteracting oxidative stress.^80–82^

Consistent with these reported biological activities, in our study, effera^®^ significantly increased SCFA production coinciding with an increase in the expression of tight junction-associated genes ZO-1 and occludin and improved epithelial barrier integrity following LPS challenge. Microbiota-derived SCFAs are also important components of the early-life intestinal environment, although their abundance and relative distribution vary according to infant age and feeding pattern. In a cohort of 163 infants aged 3-5 months, breastfeeding status was associated with distinct fecal SCFA and intermediate-metabolite profiles. Exclusively breastfed infants had a higher relative proportion of acetate and higher lactate concentrations, despite having lower absolute concentrations of several SCFAs and BCFAs than non-exclusively breastfed infants.^83^ The increased acetate, propionate, butyrate, and valerate production observed with effera^®^ therefore provides useful mechanistic guidance for an infant-relevant study. Such a study should determine whether effera^®^ promotes a metabolic profile appropriate to early life, including the balance among acetate, lactate, propionate, and butyrate, rather than evaluating total SCFA production alone. Notably, although native hmLF, bLF and effera^®^ significantly enhanced the production of SCFA, the magnitude of the barrier protective effect was greater for effera^®^ than bLF. This finding suggests that the beneficial effect of effera^®^ extends beyond SCFA production, and microbiome-associated metabolites may additionally contribute to effera^®^’s beneficial effects.

While skim milk at 50 mg/day elicited no response, measurable responses were observed at 1000 mg/day. This finding is biologically plausible due to the composition of skim milk. In addition to casein, skim milk contains whey protein, lactose, and bioactive peptides that can serve as substrates for gut microbial fermentation. The biological response observed at 1000 mg/day could be due to the composition of the dairy matrix rather than the single bioactive component.^84–86^

The increase in SCFA production was accompanied by shifts in the microbial composition. In addition to enhancing microbial activity, effera^®^ significantly induced distinct shifts in the gut microbial composition. Maintaining microbial diversity along with an increase in the beneficial bacterial abundance is a primary indicator of a healthy and resilient gut.^87^ Enhancing gut microbiomes is important as they regulate digestion, synthesize essential vitamins, and modulate the immune system.^8–10,29^ In our study, effera^®^ significantly enhanced bacterial cell density compared to bLF. At the taxonomic level, effera^®^ induced dose-dependent increases in *Bacillota_A* and *Bacteroidota*, the two most abundant phyla in the human gut microbiome.^88^ Within the *Bacillota_A*, an increase in the members of *Oscillospiraceae* family, including *Lawsonibacter_faecis* and *Hominicoprocola_fusiformis*, contribute to the SCFA production. Members of the *Oscillospiraceae* family have been reported for their strong saccharolytic (carbohydrate-fermenting) activity and capacity to generate SCFA, notably butyrate among other metabolites, increasing mucus production and intestinal epithelial integrity.^89–91^ Within the *Bacteroidota* phylum, effera^®^ promoted abundance of various species belonging to the *Bacteroidaceae* family. Members of the bacteroid family are the most abundant commensal bacteria in the human gut and play a central role in the degradation of complex dietary polysaccharides and host-derived glycans.^92,93^ *Bacteroides_thetaiotaomicron* has been extensively studied for its ability to metabolize diverse carbohydrates, promote mucus layer maintenance and influence host immune responses.^93,94^ Similarly, *Bacteroides_uniformis* has been associated with improving metabolic health by reducing serum glucose, insulin, triglycerides, and cholesterol levels in high-fat diet-fed mice.^95^ Another in vivo study reported that *Bacteroides_uniformis* has anti-inflammatory activity by mitigating diet-induced endotoxemia, and maintains intestinal homeostasis by increasing regulatory T cells and intraepithelial lymphocytes in the lamina propria.^96^ In other systems, increasing regulatory T cells can suppress inflammation and increase tolerance.^97^ effera^®^ also increased the abundance of *Rikenellaceae* family members, including *Alistipes* species. *Alistipes* species has been reported to ferment complex carbohydrates and amino acids to produce SCFA like acetate, propionate and succinate.^91^ Certain species of *Alistipes* have been shown to alleviate intestinal inflammation by regulating host lipidomes and reducing pro-inflammatory cytokines.^98,99^ In addition, abundance of *Parabacteroides_merdae* and *Parabacteroides_distasonis* was observed. Notably, *Parabacteroides_distasonis* was studied for its anti-inflammatory properties and improved intestinal barrier function. Studies have reported that it downregulates inflammatory NF-kB signaling and increases regulatory T cells. It also generates indole derivatives such as indole acrylic acid, which is known to play a vital role in mucosal repair and barrier integrity by binding to aryl hydrocarbon receptors.^100^ Notably, several bacterial taxa that were positively modulated following effera^®^ exposure in the SIFR^®^ platform were also observed to increase in fecal microbiome analyses from a human clinical study evaluating effera^®^ supplementation. These included *Lawsonibacter* within the *Oscillospiraceae* family, *Alistipes* within the *Rikenellaceae* family, and Prevotella within the *Prevotellaceae* family, providing cross-model evidence for reproducible modulation of specific microbial taxa in response to effera^®^ exposure.^49^ The consistency of these findings across controlled ex vivo fermentation and human clinical settings further supports the potential of effera^®^ to beneficially influence the composition of the gut microbiome.

The potential relevance of lactoferrin to early-life microbial development is also supported by infant studies. In term and preterm mother-infant pairs, fecal lactoferrin concentrations were positively associated with fecal *bifidobacteria* and *lactobacilli* during the first month of life, supporting a relationship between lactoferrin exposure and the establishment of the neonatal microbial environment.^101^ In addition, infants receiving a formula containing bovine milk-fat-globule membrane and bovine lactoferrin showed subtle differences in stool microbiome and metabolome profiles at 4 months of age, including an increased prevalence of selected *Bacteroides* species.^102^ These clinical observations are broadly consistent with the present finding that lactoferrin-containing treatments can modulate microbial composition and metabolic activity. Because the infant formula intervention contained both milk-fat globule membrane and lactoferrin, and because infant communities differ from adult microbiota, the present results should be viewed as providing mechanistic guidance for a dedicated infant microbiome study rather than predicting an identical taxonomic response.

### Differential microbiome and barrier responses to effera^®^ human lactoferrin vs. bovine lactoferrin

A noteworthy finding of our study demonstrates that effera^®^ outperformed dose-matched bLF across multiple endpoints including microbial cell density, microbial composition, and epithelial barrier integrity. Although both proteins exhibited microbial shifts and increases in epithelial barrier integrity, the magnitude of the response elicited by effera^®^ was greater, suggesting that effera^®^ might be interacting with the gut microbiota differently than bLF. While the underlying mechanism of action remains to be explored, several potential hypotheses may explain effera^®^’s performance. Species-specific differences in primary sequence may influence microbial utilization and metabolite generation. Bovine LF shares only 69% sequence similarity with hmLF.^103^ effera^®^ also displays a similar digestive profile to hmLF, and generates bioactive peptides more similar to hmLF.^104^ This similarly could lead to the generation of similar downstream bioactive metabolites and thus better mimic the effects of hmLF.

effera^®^ may promote abundance or activity of certain microbial populations that efficiently produce metabolites that support barrier integrity. Consistent with this hypothesis, we observed that effera^®^ induced shifts at a broader species level than bLF, enriching several taxa associated with carbohydrate fermentation and intestinal health. These compositional changes may have enhanced microbial cross-feeding, resulting in the production of various metabolites influencing intestinal health. Importantly, differences in gut barrier integrity results between effera^®^ and bLF suggest that additional fermentation-derived metabolites or microbiome-mediated signaling pathways may also contribute to the effect. This finding that effera^®^ had more pronounced gut barrier and microbial growth effects highlights the potential importance of protein origin and structure for host-microbiome interactions. However, future studies are required to identify specific metabolites and microbial pathways that may further explain these differences.

## Limitations and Future Directions

The ex vivo SIFR^®^ technology platform provides a highly controlled and reproducible framework for evaluating microbiome-mediated responses across independent human donors. Nevertheless, as will all ex vivo systems, it does not reproduce every component of whole-body physiology. Epithelial barrier function was assessed using in vitro cell culture systems which, while widely accepted for evaluating intestinal permeability, lack the ability to fully reproduce the multicellular complexity of the intestinal mucosa.^105^ While the Caco-2/THP-1 co-culture model incorporates epithelial and immune cross-talk, it does not fully capture the complexity of the intestinal environment which includes diverse immune cell populations,^106,107^ goblet cells, Paneth cells, vascularization and surrounding stromal tissues and these host cell lines are originally derived from cancer patients rather than healthy subjects.^108–110^ The experimental system also does not account for repeated intake of food and dietary compounds, changes in nutrient availability which could influence the long-term adaptation of the gut microbiome.^111,112^ Furthermore, systemic processes, including endocrine hormonal signaling and physiological body responses are also not represented in this platform.^113,114^ In addition, the modest number of donors (n = 6), which, given the substantial interindividual variability characterizing the human gut microbiome, may limit the generalizability of the ex vivo findings to the broader adult population. Future studies that incorporate a larger number of donors is suggested to strengthen the validity of the observed findings. Thus, we interpret the observed effects on barrier integrity as evidence of biological activity rather than a direct confirmation of clinical efficacy.

Our ex vivo modeling was limited to comparisons between effera^®^ and bLF and did not include native hmLF as a reference control across all doses. effera^®^ was evaluated for a broad dose range, while bLF was evaluated at only 3 dose levels. As a result, the full dose-response relationship was established but with limited direct comparisons between the two proteins at certain equivalent doses. Finally, although significant changes in microbial composition, barrier integrity and immune responses were observed, the present study does not establish a direct causal relationship between individual metabolites and epithelial responses. The specific microbial metabolites other than SCFA were not identified and require further investigation. Multiple fermentation-derived products may contribute collectively to the observed biological effects.

## Novelty and Significance

Previous studies investigating gut health effects of LF have largely focused on direct exposure of intact LF in epithelial or immune cell models, or on its ability to influence the growth of individual bacterial species.^43,115–122^ While these studies have established important antimicrobial, immunomodulatory, and probiotic-supporting properties of LF, they do not fully capture the complex sequence of events following oral consumption, where LF undergoes digestion, and extensive interaction with the gut microbiome before interacting with the intestinal epithelium. The present study addresses this knowledge gap by evaluating recombinant hLF within an integrated microbiome-host framework that connects microbial fermentation and compositional changes with downstream epithelial-barrier and immune outcomes.

A central novelty of this study is the side-by-side evaluation of intact and colonic fermentation-derived LF preparations within the same experimental system. This design revealed that the strongest barrier-supporting activity emerged following microbial fermentation rather than following exposure to the intact protein alone, highlighting the critical role of the gut microbiota. Importantly, this conclusion was supported across multiple complementary readouts, including SCFA production, bacterial growth and composition, and gut barrier enhancement via TEER. By linking LF biotransformation to changes in microbial metabolism, epithelial resistance, tight junction-associated gene expression, and inflammatory signaling, the study identifies microbial fermentation as an important contributor to the functional activity of LF. Thus, rather than examining the microbiome and host response as separate endpoints, this workflow connects them as sequential and biologically interdependent processes.

An additional strength of this work is the direct comparison of hLF and bLF under identical digestion, fermentation, and host-response conditions. Although both proteins stimulated microbial activity, effera^®^ demonstrated greater responses across several epithelial-barrier readouts following microbial fermentation. This head-to-head design provides new evidence that species-specific differences in LF protein structure and gastrointestinal processing may influence microbial utilization and downstream biological activity. The findings therefore extend beyond demonstrating LF functionality and provide a mechanistic basis for investigating whether hLF offers distinct functional advantages within the human gut ecosystem.

## Conclusion

In summary, this study demonstrates that the gut-related activity of LF is strongly shaped by its interactions with the gut microbiome. Microbial fermentation emerged as a key step in translating LF into its downstream effects. Fermentation-derived products of effera^®^ increased SCFA production, strengthened TEER under both basal and LPS-challenged conditions, upregulated the tight -junction-associated targets ZO-1 and occludin, and reduced CXCL-10 and IL-8 responses. Across several functional readouts, effera^®^ produced greater responses than bLF supporting the concept that hLF may possess distinct functional properties within the human gut ecosystem. Together, these findings provide a mechanistic framework linking microbiome-mediated biotransformation of LF to improved intestinal barrier function and support the future development of how human-equivalent bioactive proteins can be leveraged to support intestinal health.

